# The Prevalence of Inappropriate Image Duplication in Biomedical Research Publications

**DOI:** 10.1101/049452

**Authors:** Elisabeth M. Bik, Arturo Casadevall, Ferric C. Fang

**Affiliations:** Department of Medicine, Division of Infectious Diseases, Stanford School of Medicine, Stanford, California, USA; Department of Molecular Microbiology and Immunology, Johns Hopkins Bloomberg School of Public Health, Baltimore, Maryland, USA; Department of Medicine, Johns Hopkins School of Medicine, Baltimore, Maryland, USA; Departments of Laboratory Medicine and Microbiology, University of Washington School of Medicine, Seattle, Washington, USA

**Keywords:** Research misconduct, ethics in science, biomedical research, peer review

## Abstract

Inaccurate data in scientific papers can result from honest error or intentional falsification. This study attempted to determine the percentage of published papers containing inappropriate image duplication, a specific type of inaccurate data. The images from a total of 20,621 papers in 40 scientific journals from 1995-2014 were visually screened. Overall, 3.8% of published papers contained problematic figures, with at least half exhibiting features suggestive of deliberate manipulation. The prevalence of papers with problematic images rose markedly during the past decade. Additional papers written by authors of papers with problematic images had an increased likelihood of containing problematic images as well. As this analysis focused only on one type of data, it is likely that the actual prevalence of inaccurate data in the published literature is higher. The marked variation in the frequency of problematic images among journals suggest that journal practices, such as pre-publication image screening, influence the quality of the scientific literature.

## IMPORTANCE

The scientific literature is cumulative, and the reproducibility of biomedical research is a topic of increasing concern. Inaccurate data, whether resulting from honest error or intentional misconduct, can contribute to research irreproducibility, but the prevalence of such data is unknown. Here, we provide the first estimate of the percentage of a specific type of inaccurate data, inappropriate image duplication, in published biomedical research papers. Approximately 1 of every 25 papers was found to contain some form of problematic image, with a substantial increase in prevalence since 2003. Current standards appear insufficient to prevent flawed papers from being published. Greater efforts are needed to ensure the reliability and integrity of the research literature.

## INTRODUCTION

Inaccuracies in scientific papers have many causes. Some result from honest mistakes, such as incorrect calculations, use of the wrong reagent, or improper methodology (1). Others are intentional and constitute research misconduct, including situations in which data are altered, omitted, manufactured or misrepresented in a way that fits a desired outcome. The prevalence of honest error and misconduct in the scientific literature is unknown. One review estimated the overall frequency of serious research misconduct, including plagiarism, to be 1% (2). A meta-analysis by Fanelli, combining the results of 18 published surveys, found that 1.9% of researchers have admitted to modification, falsification or fabrication of data (3).

There is also little firm information on temporal trends regarding the prevalence of error and misconduct. Research error and misconduct have probably always existed. Even scientific luminaries such as Darwin, Mendel, and Pasteur have been accused of manipulating or misreporting their data (4, 5). However, the perception of error and misconduct in science has been recently magnified by high profile cases and a sharp rise in the number of retracted manuscripts (6). In recent years, retractions have increased at a rate that is disproportionately greater than the growth of the scientific literature (7). Although this could be interpreted as an increase in problematic papers, the actual causes may be more complex and could include a greater inclination by journals and authors to retract flawed work (7). Retractions are a poor indicator of error because most retractions result from misconduct (8), and many erroneous studies are never retracted (1). In fact, only a very small fraction of the scientific literature has been retracted. As of April 2016, the PubMed bibliographic database listed 8,735 retracted publications among more than 25 million articles (0.035%).

Concerns about misconduct have been accompanied by increasing concerns about the reproducibility of the scientific literature. An analysis of 53 landmark papers in oncology reported that only 6 could be reproduced (9), and other pharmaceutical industry scientists have also reported low rates of reproducibility of published findings, which in some cases led to the termination of drug development projects (10). In the field of psychology, less than half of experimental and correlational studies are reportedly reproducible (11). Inaccurate data can result in societal injury. For example, a now-retracted study associating measles vaccination with autism continues to resonate and may be contributing to low vaccination rates (12). Corrosion of the literature, whether by error or misconduct, may also impede the progress of science and medicine. For example, false leads may be contributing to increasing disparities between scientific investment and measurable outcomes, such as the discovery of new pharmacological agents (13).

In this study we sought to estimate the prevalence of a specific type of inaccurate data that can be readily observed in the published literature, namely inappropriate image duplication. The results demonstrate that problematic images are disturbingly common in the biomedical literature and may be found in approximately 1 out of every 25 published articles containing photographic image data, in particular Western blots.

## RESULTS

### Papers containing inappropriately duplicated images

A total of 20,621 research papers containing the search term “western blot” from 40 different journals and 14 publishers were examined for inappropriate duplications of photographic images, with or without repositioning or evidence of alteration (Table S1). Of these, 8,138 (39.8%) were published by a single journal (*PLOS ONE*) in 2013 and 2014; the other 12,483 (60.5%) papers were published in 39 journals spanning the years 1995-2014 (Fig. 1). Overall, 782 (3.8%) of these papers were found to include at least one figure containing inappropriate duplications.

**Figure 1.**
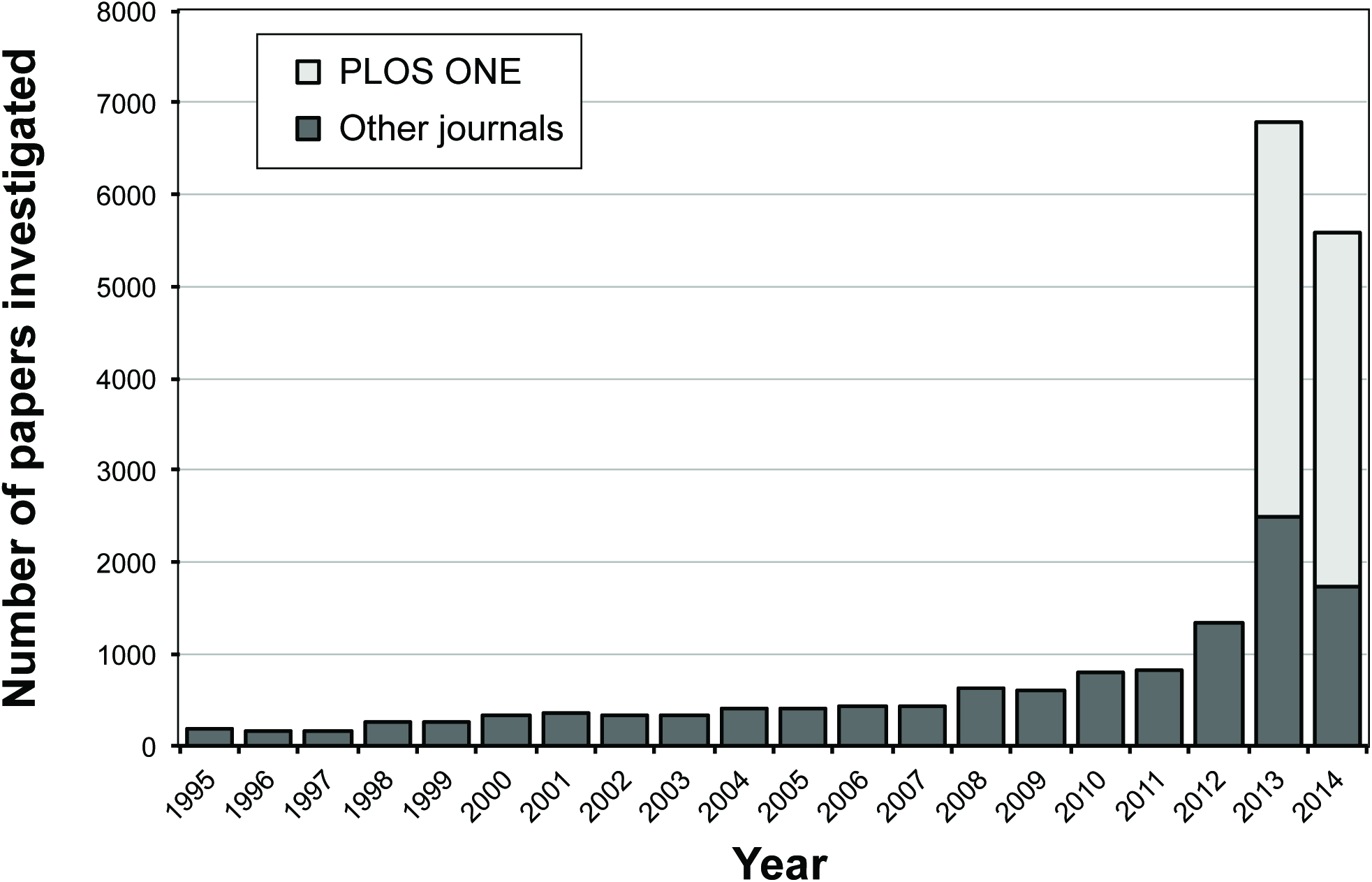
Publications investigated in this study by year of publication. The majority of screened papers were published in 2013 and 2014 due to the large proportion of *PLOS ONE* papers (39.5%) in the dataset. The lowest number of papers (n=151) screened in this study was published in 1996.

### Classification of inappropriately duplicated images

Problematic images were classified into three major categories: simple duplications, duplications with repositioning, and duplications with alteration.

- Category I: Simple Duplications. Figures containing two or more identical panels, either within the same figure or between different figures within the same paper, purporting to represent different experimental conditions, were classified as simple duplications. The most common examples in this category were beta-actin loading controls that were used multiple times to represent different experiments or identical microscopy images purporting to be obtained from different experiments. For papers containing such figures, the methods and results were reviewed to establish that the duplicated figures were indeed re-used for different experiments. The re-use of loading controls in different figures obtained from the same experiment was not considered to be a problem. Examples of simple duplication are shown in Fig. 2.
- Category II: Duplication with Repositioning. This category included microscopic or blot images with a clear region of overlap, where one image had been shifted, rotated, or reversed in respect to the other. Fig. 3 shows examples of duplicated figures with repositioning.
- Category III: Duplication with Alteration. This category consisted of images that were altered with complete or partial duplication of lanes, bands, or groups of cells, sometimes with rotation or reversal in respect to each other, within the same image panel or between panels or figures. This category also includes figures containing evidence of “stamping” in which a defined area is duplicated multiple times within the same image, “patching” in which part of an image is obscured by a rectangular area of different background, and FACS images sharing conserved regions and other regions in which some data points have been added or removed. Examples of duplicated images with alteration are shown in Fig. 4.

**Figure 2.**
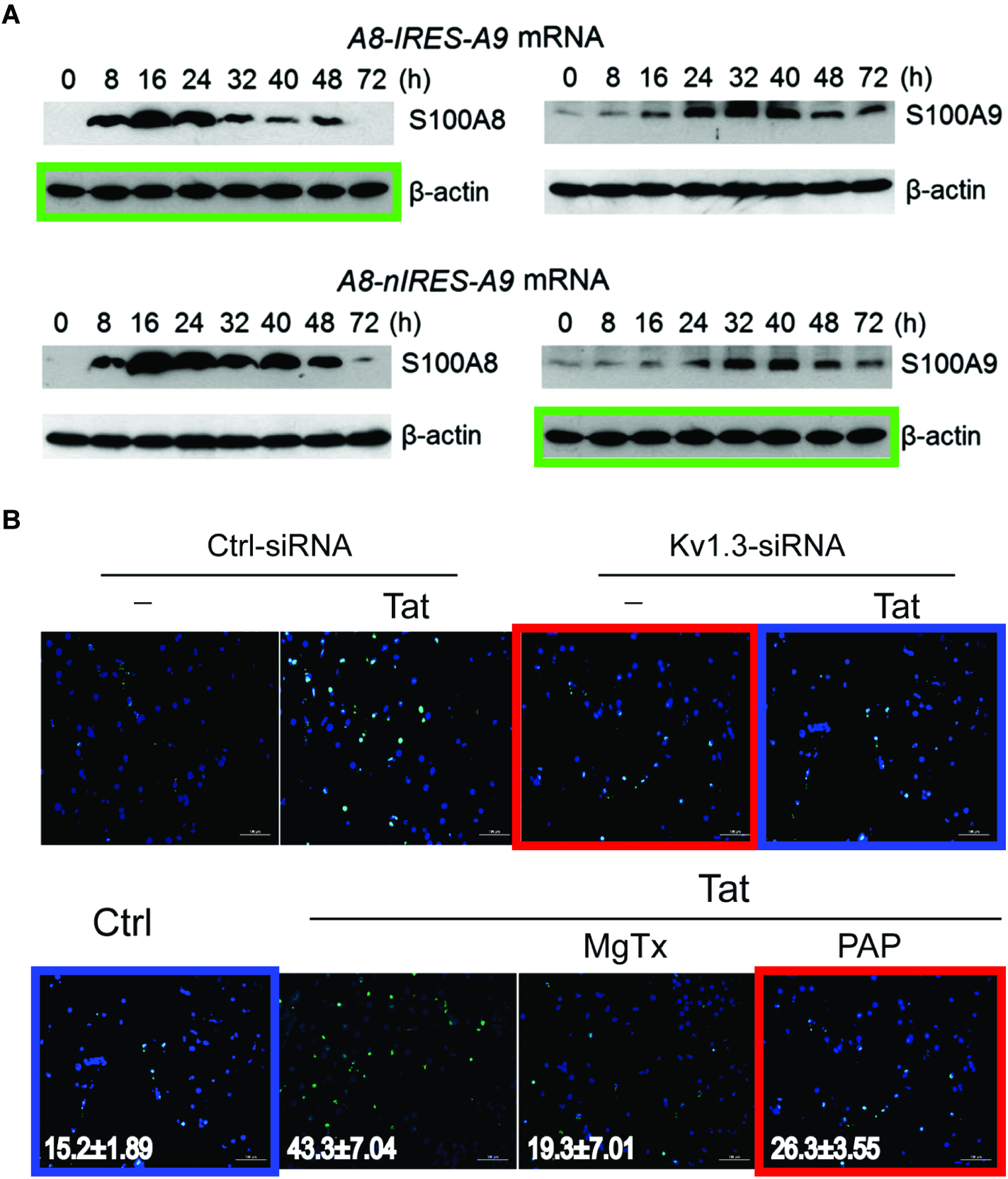
Examples of simple duplication (Category I). **A**. The beta-actin control panel in the top left is identical to the panel in the bottom right (shown by green boxes), although each panel represents a different experimental condition (27); corrected in: (28). Figure reproduced with permission from the publisher. **B**. The panels shown here were derived from two different figures within the same paper. Two of the top panels appear identical to two of the bottom panels, but they represent different experimental condition (shown with red and blue boxes) (29); corrected in: (30). Figure reproduced under the Creative Commons (CC BY) license. All duplications might have been caused by honest errors during assembly of the figures.

**Figure 3.**
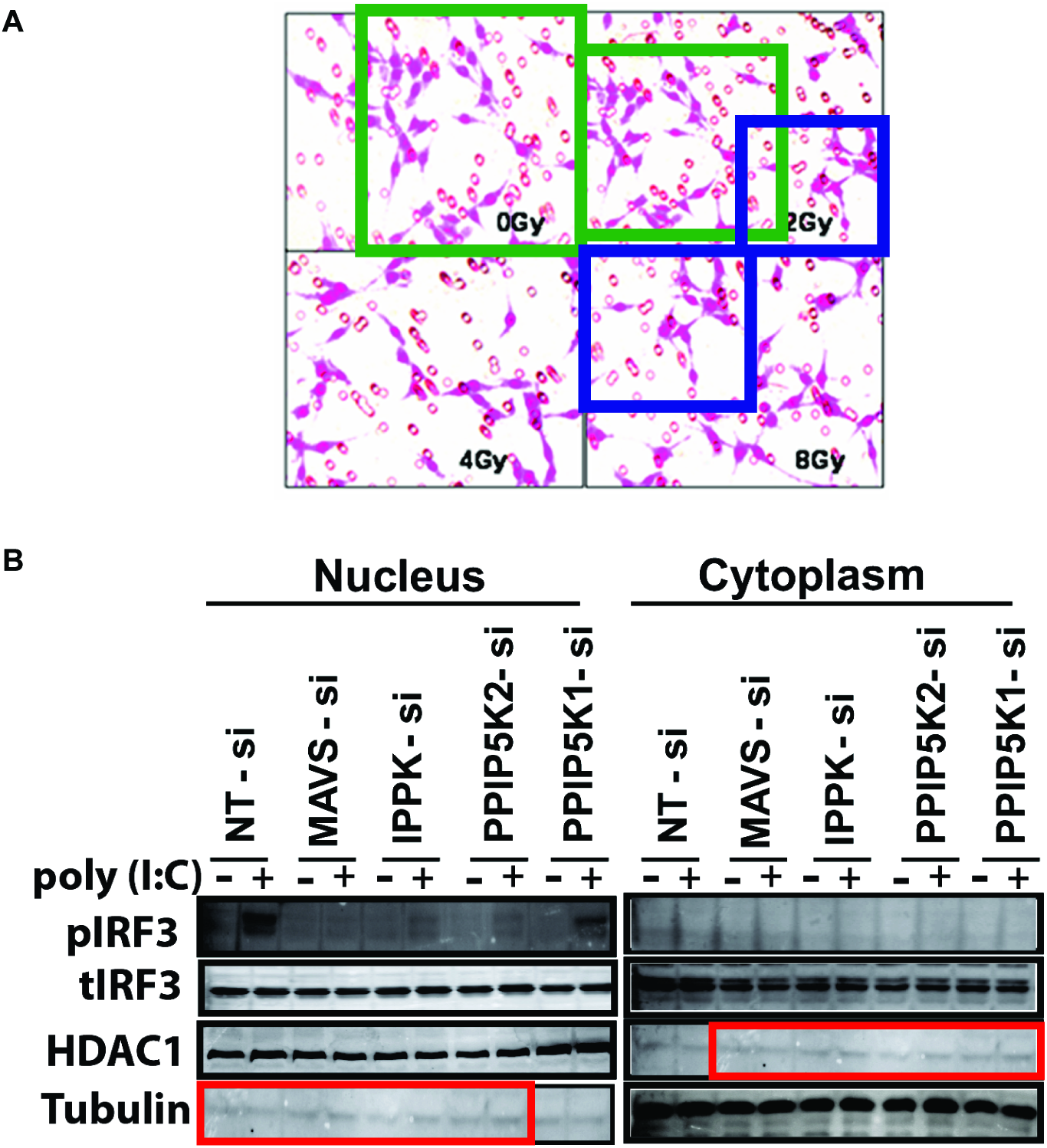
Examples of duplication with repositioning (Category II). **A**. Although the panels represent four different experimental conditions, three of the four panels appear to show a region of overlap (shown with green and blue boxes), suggesting that these photographs were actually obtained from the same specimen (31); corrected in: (32). **B**. Western blot panels “Nucleus-Protein D” and “Cytoplasm Protein C” purportedly depict different proteins and cellular fractions, but the blots appear very similar albeit shifted by two lanes (shown with red boxes) (33); corrected in: (34). Both figures reproduced under the Creative Commons (CC BY) license.

**Figure 4.**
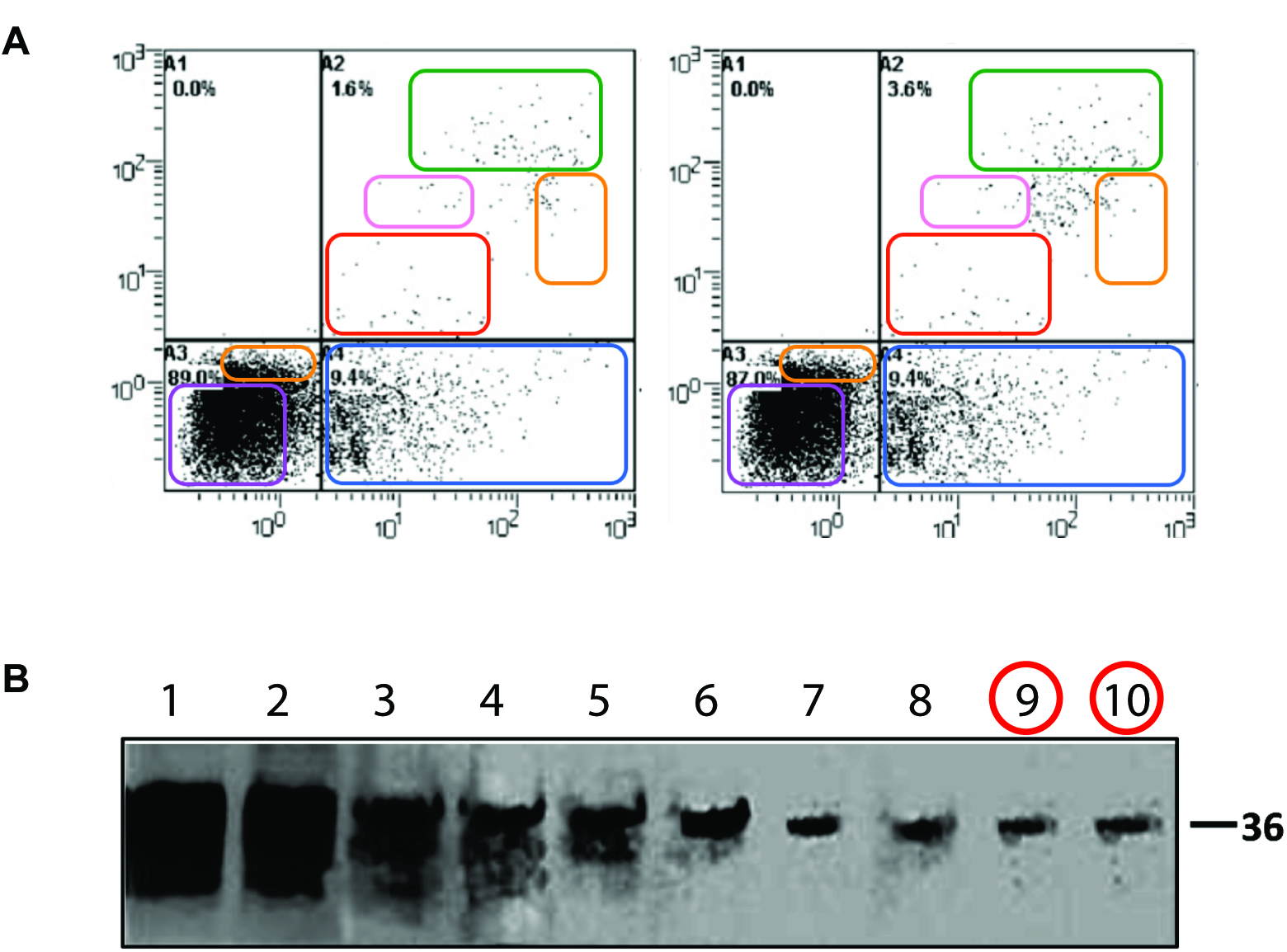
Examples of duplication with alteration (Category III). **A**. The left and right FACS panels represent different experimental conditions and show different percentages of cell subsets, but regions of identity (colored boxes) between the panels suggest that the images have been altered (35); retracted in (36). Figure reproduced with permission from the publisher. **B**. The figure shown here displays a Western blot of 10 different protein fractions isolated from a density gradient. The figure appears to show a single blot, but the last two lanes (highlighted with red circles) appear to contain an identical band. Exposure was altered to bring out details (37); corrected in: (38). Figure reproduced under the Creative Commons (CC BY) license.

Two additional types of image modification were not scored as problematic, although they may represent questionable research practices by current standards and would not be accepted by certain journals (*i.e., Journal of Cell Biology*):

- Cuts. Abrupt vertical changes in the background signal between adjacent lanes in a blot or gel suggest that the lanes were not next to each other in the original gel. These splices are of potential concern but do not necessarily indicate inaccurate data representation.
- Beautification. Part of the background of a blot or gel where no band of interest is expected may show signs of patching, perhaps to remove a smudge or stain. This is not considered to represent best practice according to contemporary guidelines for data presentation (14) but does not necessarily indicate inaccurate data representation.

Although researcher intent could not be definitively determined in this study, the three categories of duplicated images were felt to have different implications with regard to the likelihood of scientific misconduct. Category I (simple duplication) images are most likely to result from honest errors, in which an author intended to insert two similar images but mistakenly inserted the same image twice. Alternatively, simple duplications may result from misconduct, for example, if an author intentionally recycled a control panel from a different experiment because the actual control was not performed. Category II (duplication with repositioning) and category III (duplication with alteration) may be somewhat more likely to result from misconduct, as conscious effort would be required for these actions.

In our study, a paper was classified as containing an inappropriate duplication when at least one category I, II, or III problem was identified. Papers were classified according to the highest category of duplicated image; e.g., a paper containing both category I and category II images was classified as a category II paper.

Among the 782 problematic papers found in this study, 230 (29.4%) contained simple duplications, 356 (45.5%) contained duplicated images with repositioning, while the remaining 196 (25.1%) contained duplicated figures with alteration.

### Temporal trends in image duplication

To investigate the prevalence of image duplications and alterations over time, we plotted the percentage of papers containing inappropriate image duplication as a function of publication year (Fig. 5). The percentage of papers with image duplications appeared to be relatively low (< 2%) from 1995-2002, with no problematic images found among the 194 papers screened from 1995. However, a sharp increase in the percentage of papers with duplicated images was observed in the year 2003 (3.6%), after which the percentages have remained close to or above 4%. This pattern remained very similar when only a subset of 16 journals for which papers were scanned from all 20 years was considered, except for a decline in the duplications found in 2014, the last year of our screen (2.2%) (Fig. 5).

**Figure5.**
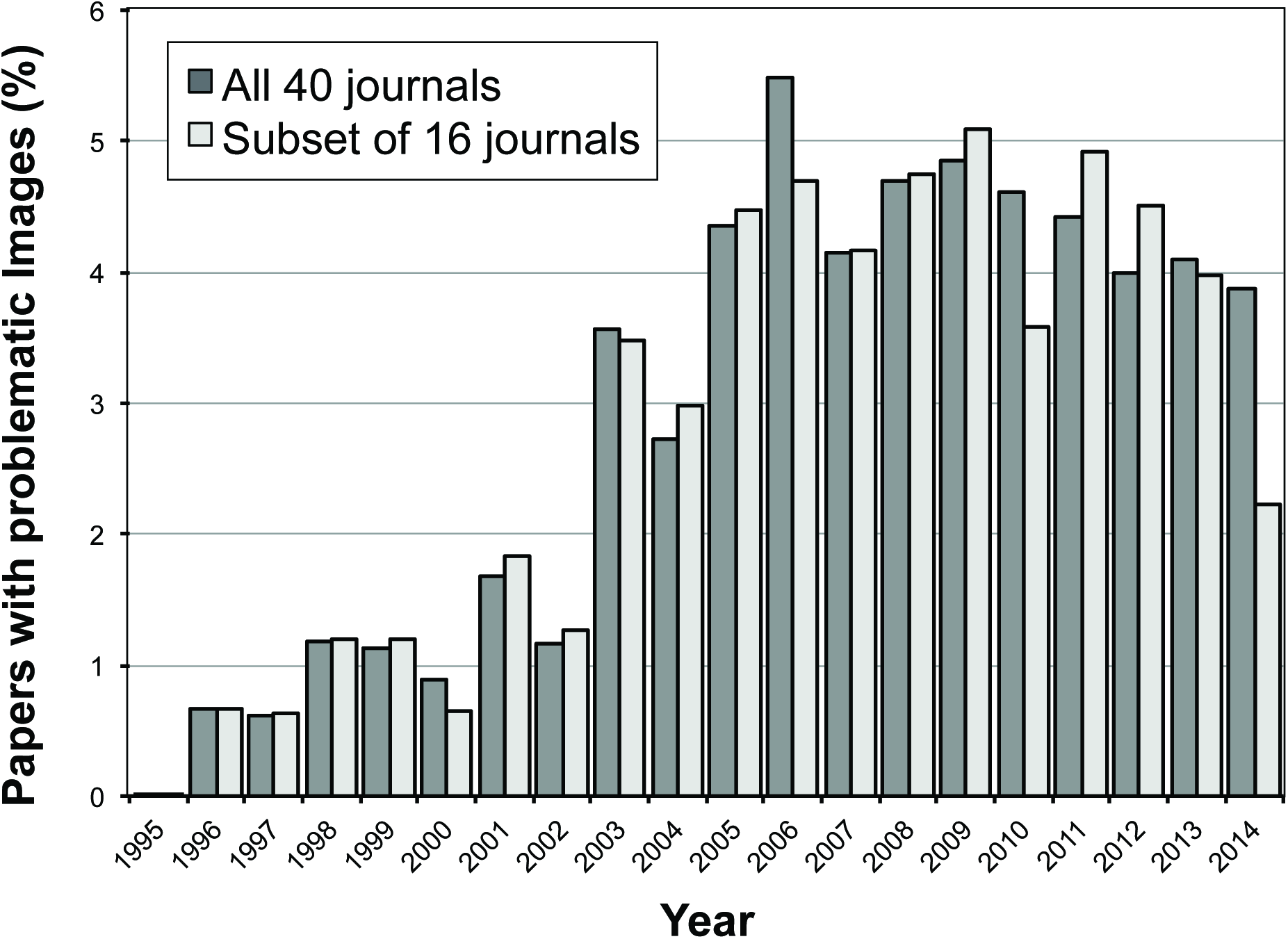
Percentage of papers containing inappropriate image duplications by year of publication. No papers with duplications were found in 1995. The dark gray bars show the data for all 40 journals. The light gray colored bars show a subset of 16 journals for which papers spanning the complete timespan of 20 years were scanned. The total numbers of papers screened in each year are shown in Figure 1.

### Correlation of impact factor with image duplication

Substantial variation in the prevalence of papers with image duplication was observed among the 40 journals investigated. In *PLOS ONE*, from which the largest number of papers was screened, 4.3% of the papers were found to contain inappropriately duplicated images, whereas the percentage of papers with image duplication ranged from 0.3% (*Journal of Cell Biology*) to 12.4% (*International Journal of Oncology*) among the other journals, with a mean of 4.4% in the last decade. Hence, even though *PLOS ONE* was the journal that provided the largest set of papers evaluated in this study, it is not an outlier with regard to inappropriately duplicated images relative to the other journals examined. To assess the possibility that journals with higher impact factors might be better at detecting problematic images and/or authors might be more careful in preparing images for publication in such journals, the relationship between the prevalence of image duplication and journal impact factor was examined (Fig. 6). For this analysis, only papers published between 2005 and 2014 were included, because the prevalence of problematic images was lower in older publications, and older papers were only evaluated for selected journals. A negative correlation between image duplication and journal impact factor was observed (Pearson’s correlation; p-value 0.019), with the lowest percentage of problematic images found in journals with high impact factors. The prevalence of image duplication in 12 open access journals was not significantly different from that in 28 non-open access journals (p = 0.38, chi-squared test).

**Figure 6.**
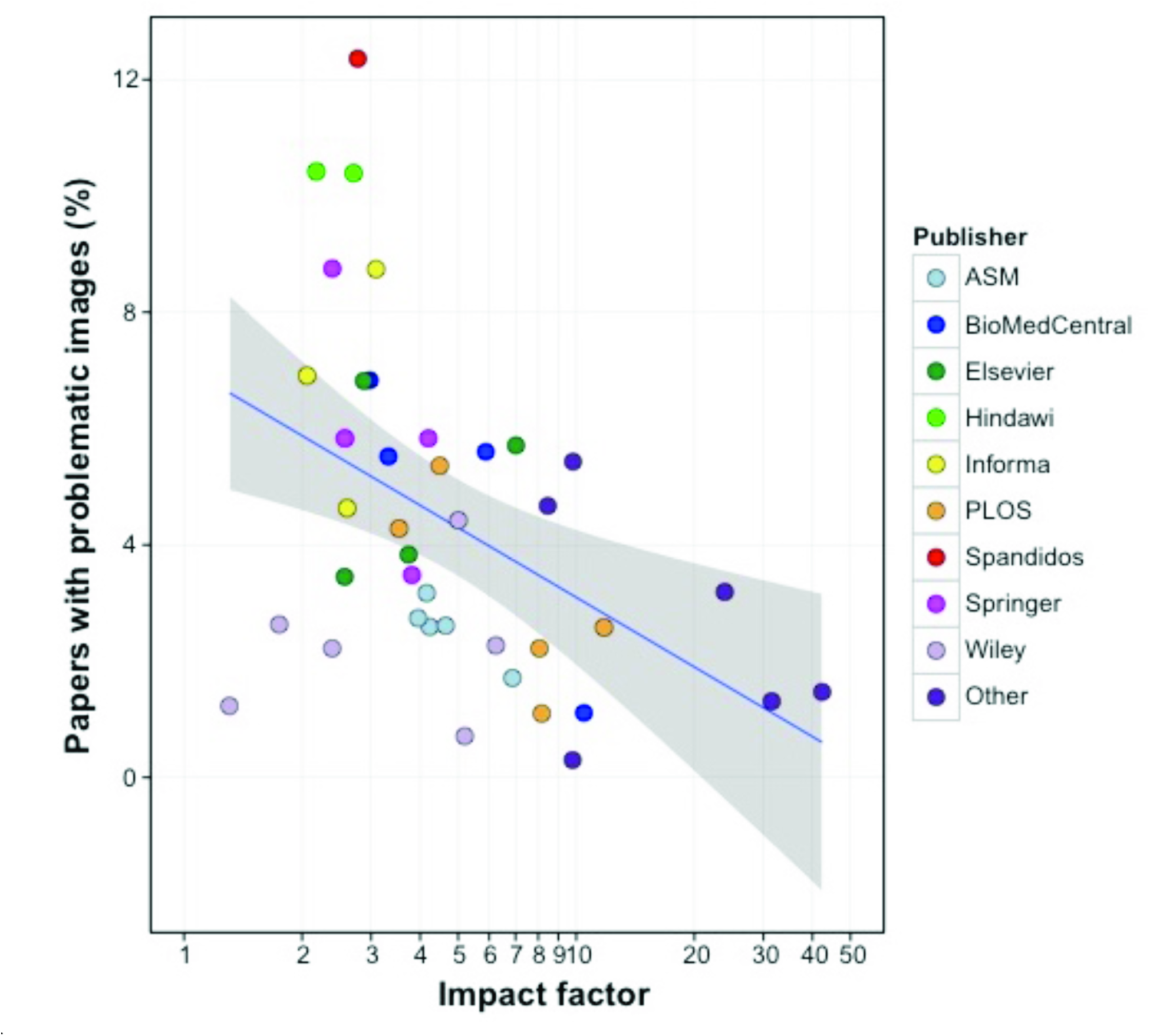
Correlation between journal impact factor and percentage of papers with image duplication. Only papers from 2005-2014 (n = 17,816) were included in this analysis. Each data point represents a journal included in this study (n=40), with data points color-coded according to their publisher (n=14; journals published by AAAS, Nature, Cell Press, the National Academy of Sciences, and the Rockefeller University Press are grouped under “Other”). The x-axis is shown on a logarithmic scale due to the small number of journals with a high impact factor included in this study. The blue line shows a linear regression model. The grey zone depicts the 95% confidence interval.

### Country of origin of papers containing image duplication

To determine whether inappropriate image duplication was more frequent in some countries than in others, the country of origin for each of the 348 papers from *PLOS ONE* containing duplicated images was compared to the country of origin for all papers published by that journal during the same time interval that were included in our search. In cases where the authors of a paper were affiliated with institutions in multiple countries, all countries were taken into account. A majority of the 8,138 screened papers published in *PLOS ONE* during the 16-month study period from 2013-2014 were affiliated with China (26.2%) and the US (40.9%) (Fig. 7). However, papers from China had a 1.89-fold higher probability of containing problematic images than would have been predicted from the frequency of publication (chi-squared test, p-value < 0.001), while papers from the US had a lower probability (0.60-fold) (chi-squared test, p-value < 0.001). Other countries with a higher-than-predicted ratio of papers containing image duplication were India (1.93) and Taiwan (1.20), whereas the prevalence of image duplication was lower than predicted in papers from the UK (0.47), Japan (0.26) and Germany (0.34).

**Figure 7.**
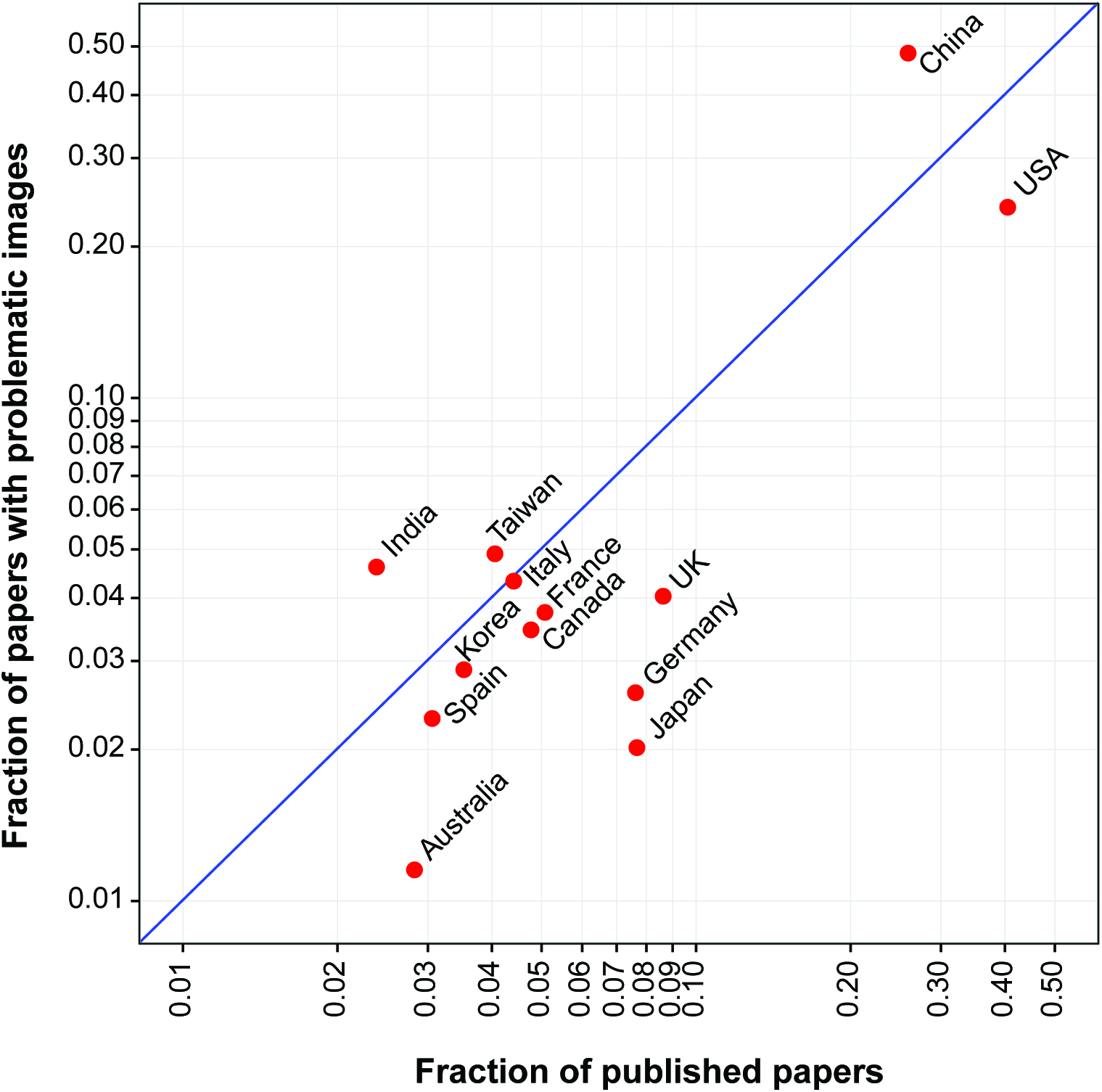
Proportion of papers with image duplications by country. The proportion of papers affiliated with specific countries submitted to *PLOS ONE* during a 16-month period in the years 2013 and 2014 (n = 8,138) is plotted versus the proportion of *PLOS ONE* papers from that same period containing inappropriate image duplication affiliated with specific countries (n = 348). Each data point represents a country for which 100 or more papers were screened. Some papers were affiliated with more than one country. The blue line represents the line where data points are expected to fall if problematic papers are distributed as expected according to their representation in the journal. Countries plotted above the blue line had a higher-than-expected proportion of problematic papers; countries plotted below the line had a lower-than-expected ratio.

### Number of authors and image duplication

Errors or misconduct might be predicted to be more frequent in papers with fewer authors, due to reduced scrutiny. The mean number of authors per paper for the 781 papers containing inappropriate image duplication was 7.28. No significant difference between the mean number of authors for papers with or without image duplication was found (p > 0.1).

### Problematic images in multiple papers by the same author

Our dataset of 782 problematic papers contained 28 papers (i.e., 14 pairs) of papers with a common first author. To determine whether authors of papers containing inappropriate image duplication were more likely to have published additional papers containing image duplication, we screened other papers written by the first and last authors of 559 papers (all from unique first authors) identified during initial screening. This analysis encompassed 2,425 papers, or a mean of 4.3 additional papers for each primary paper. In 217 cases (38.8%), at least one additional paper containing duplicated images was identified. In total, 269 additional papers containing duplicated images (11.1%) were found out of the 2,425 papers in the secondary dataset. The percentage of papers with duplicated images in the secondary dataset was significantly higher than that of the first dataset (11.1% vs. 3.8%, chi-squared test: p<0.001), indicating that other papers by first or last authors of papers with duplicated images have an increased probability of also containing duplicated images.

## DISCUSSION

The quality and integrity of the scientific literature has been increasingly questioned (8, 9, 15, 16). The present study attempts to empirically determine the prevalence of one type of problematic data, inappropriate image duplication, by visual inspection of electrophoretic, microscopic and flow cytometric data from more than 20,000 recent papers in 40 primary research journals. The major findings of this study are: (1) figures containing inappropriately duplicated images can be readily identified in published papers through visual inspection without the need for special forensic software methods or tools; (2) approximately 1 of every 25 published papers contains inappropriately duplicated images; (3) the prevalence of papers with inappropriate image duplication rose sharply after 2002 and has since remained at increased levels; (4) the prevalence of inappropriate image duplication varies among journals and correlates inversely with journal impact factor; (5) papers containing inappropriately duplicated images originated more frequently from China and India and less frequently from the US, UK, Germany, Japan, or Australia; and (6) other papers by authors of papers containing inappropriately duplicated images often contained duplicated images as well. These findings have important implications for the biomedical research enterprise and suggest a need to improve the literature though greater vigilance and education.

The finding that figures with inappropriate duplications can be readily identified by simple inspection suggests that greater scrutiny of publications by authors, reviewers, and editors might be able to identify problematic figures prior to publication. The *Journal of Cell Biology* was among the first to call attention to the problem of figure alteration in manuscripts (17), and this journal instituted a policy to carefully inspect all manuscripts for image manipulation prior to publication (14). The low prevalence of problematic images in this journal (0.3%) suggests that these measures have been effective. The *EMBO Journal*, which was not part of the present study, has also instituted a manual screening process for aberrant images (18). Our findings are consistent with the notion that greater scrutiny by journals can reduce the prevalence of problematic images. However, this is likely to require a concerted effort by all journals, so that authors of papers with problematic data do not simply avoid publication in venues that employ rigorous screening procedures.

The prevalence of papers containing inappropriate image duplication increased markedly in 2003 and has remained high in subsequent years. This coincides with the observed increase in retracted publications (8) and provides empirical evidence that the increased prevalence of problematic data is not simply a result of increased detection, as has been suggested (19). Although the causes of the increased frequency of image duplication since 2003 are not known, we have considered several possible explanations. First, older papers often contain figures with lower resolution, which may have obscured evidence of manipulation. Second, the widespread availability and usage of digital image modification software in recent years may have provided greater opportunity for both error and intentional manipulation. Third, the increasing tendency for images to be directly prepared by authors instead of by professional photographers working in consultation with authors has removed a potential mechanism of quality control. One possible mechanism to reduce errors at the laboratory level would be to involve multiple individuals in the preparation of figures for publication. The lack of correlation between author number and the frequency of image duplication suggests that the roles of most authors are compartmentalized or diluted, such that errors or misconduct are not readily detected. A fourth consideration is that increasing competition and career-related pressures may be encouraging careless or dishonest research practices (20). Finally, electronic manuscript submission, as implemented by many journals in the early 2000s, facilitated submissions from countries that were previously discouraged to submit because of high postal costs.

A large variation in the prevalence of papers containing inappropriately duplicated images was observed among journals ranging from the *Journal of Cell Biology* (0.3%) to the *International Journal of Oncology* (12.4%), a difference of more than 40-fold. The differences among journals are important because, as noted above, these suggest that journal editorial policies can have a substantial impact on this problem. Alternatively, the variable prevalence of duplication could be partly accounted for by variations in the average number of figures and panels-per-figure, which is likely to differ per journal but was not determined in our study. The inverse correlation between the prevalence of problematic papers and journal impact factor contrasts with the positive correlation observed for research misconduct resulting in retraction 8, 21–23). Although the association was weak, this may suggest that higher impact journals are better able to detect anomalous images prior to publication. Alternatively, authors submitting to such journals may be more careful with figure preparation. Nevertheless, we note that even the most highly selective journals contain some papers with figures of concern.

China and the United States were responsible for the majority of papers containing inappropriately duplicated images, which is not surprising given the large research output of these countries. However it is noteworthy that the proportion of *PLOS ONE* papers from China and India that were found to contain problematic images was higher than would be predicted from their overall representation, whereas the opposite was true for papers from the US, UK, Germany, Japan and Australia. This suggests that ongoing efforts at scientific ethics reform in China and India should pay particular attention to issues relating to figure preparation (24, 25). The analysis of geographic origin was limited to papers published in *PLOS ONE*, because this journal offered an online tool to search for this information. The geographic distribution of papers with problematic images may be different in other journals.

In nearly 40% of the instances in which a problematic paper was identified, screening of other papers from the same authors revealed additional problematic papers in the literature. This suggests that image duplication results from systematic problems in figure preparation by individual researchers, which tend to recur.

Our findings suggest that as many as 1 out of every 25 published papers containing Western blot or other photographic images could contain data anomalies. This is likely to be an underestimate of the extent of problematic data in the literature for several reasons. First, only image data were analyzed, thus errors or manipulation involving numerical data in graphs or tables would not have been detected. Second, only duplicated images within the same paper were examined, thus the reuse of images in other papers by the same author(s) would not have been detected. Third, since problematic images were detected by visual inspection, the false-negative rate could not be determined; we readily acknowledge that our screen may have missed problematic papers. It should be noted that our findings contrast with a recent small study by Oksvold that examined 120 papers from 3 different cancer research journals, reporting duplicated images in 24.2% of the papers examined (26). Many of the reported image duplications in the Oksvold study involved representation of identical experiments, which we do not regard as necessarily inappropriate, as this form of duplication does not alter the research results. For comparison, we screened 427 papers from the same 3 journals examined by Oksvold and found the average percentage of problematic papers in these journals to be 6.8, which is closer to our findings for other journals. Moreover, our study included more than 20,000 papers from 40 journals; in addition to more rigorous inclusion criteria, we required consensus between three independent examiners for an image to be classified as containing inappropriate duplication, ensuring a low false-positive rate.

The high prevalence of inaccurate data in the literature should be a finding of tremendous concern to the scientific community, since the literature is the record of scientific output upon which future research progress depends. Papers containing inaccurate data can reduce the efficiency of the scientific enterprise by directing investigators to pursue false leads or construct unsupportable hypotheses. Although our findings are disturbing, they also suggest specific actions that can be taken to improve the literature. Increased awareness of recurring problems with figure preparation, such as control band duplication, can lead to the reform of laboratory procedures to detect and correct such issues prior to manuscript submission. The variation among journals in the prevalence of problematic papers suggests that individual journal practices, such as image screening, can reduce the prevalence of problematic images (17, 18). The problems identified in this study provide further evidence for the scientific establishment that current standards are insufficient to prevent flawed papers from being published. Our findings call for the need of greater efforts to ensure the reliability and integrity of the research literature.

## MATERIALS AND METHODS

### Selection strategy

A total of 20,621 papers were selected from 40 different scientific journals in the fields of microbiology and immunology, cancer biology, and general biology. These journals were published by 14 organizations (average of 2.9 journals per publisher, range 1-6) (Table S1). All journals included in the search were indexed in PubMed, with a mean impact factor of 6.9 (range 1.3 - 42.4, Thomson Reuters 2013). Papers were examined if they contained the search term “western blot”, using the search tool provided at the journal’s website. Only original research papers containing figures were included; retracted papers, review papers, and conference abstracts were excluded. Corrected papers were included only if the correction involved issues other than the problems identified by the present analysis (e.g., incorrect grant statement). From a single journal (*PLOS ONE)*, 8,138 papers published in 2013 and 2014 were included in the study, comprising 39.5% of the dataset. From the remaining journals, a mean of 320.1 (range 77 - 1070) papers per journal were included. For most of these journals, if more than 50 papers were found in a given year, screening was limited to the first 40-50 papers that were shown in the search field. The selected papers spanned the years 1995-2014, with most papers published in 2013-2014, primarily as a result of the large contribution of papers from *PLOS ONE* (Fig. 1). The large number of *PLOS ONE* papers analyzed reflects both the journal format, which facilitates image analysis, and the fact that *PLOS ONE* is currently the world’s largest scientific journal with approximately 30,000 new articles per year (https://en.wikipedia.org/wiki/PLOS_ONE).

### Visual screening

All papers were screened by examining images at the publisher’s website. Although papers were selected using the search term “western blot”, all types of photographic image were examined, including protein and nucleic acid electrophoretic gels and blots, histology or microscopy images, and photos of culture plates, plants, and animals. Fluorescence-activated cell sorting (FACS) plots were included as well since these, like photographic images, purportedly represent raw data. Figure panels containing line art such as bar graphs or line graphs were not included in the study. Images within the same paper were visually inspected for inappropriate duplications, repositioning, or possible manipulation (e.g., duplications of bands within the same blot). All papers were initially screened by one of the authors (EMB). If a possible problematic image or set of images was detected, figures were further examined for evidence of image duplication or manipulation using the Adjust Color tool in Preview software on an Apple iMac computer. No additional special imaging software was used. Supplementary figures were not part of the initial search but were examined in papers in which problems were found in images in the primary manuscript. All figures found by the screening author (EMB) to contain possible duplications were independently reviewed by the two co-authors (FCF and AC). Consensus had to be reached among all three authors for a paper to be considered to contain unequivocal evidence of inappropriate figure duplication. Consensus among all three authors was reached in 90.4% of the papers selected during primary screening.

### Statistical analysis

The R software package was used to plot and analyze data. The stat_smooth method implemented in the R ggplot2 library was applied to find the best fit for a linear regression model on semi-log transformed data examining the relationship between the 2013 Thomson Reuters impact factor and percentage of papers with inappropriately duplicated images for the 40 journals included in this study. Pearson’s correlation and Pearson’s Chi-square tests with Yates’ continuity correction were performed in basic R. R code is available as Supplemental Data S1.

### FUNDING INFORMATION

This research received no specific grant from any funding agency in the public, commercial, or not-for-profit sectors.

## SUPPLEMENTAL MATERIAL

1. Supplemental Table S1: The 40 journals screened in this study

2. Supplemental Data S1: R code

## SUPPLEMENTARY TABLE

**Table S1.**
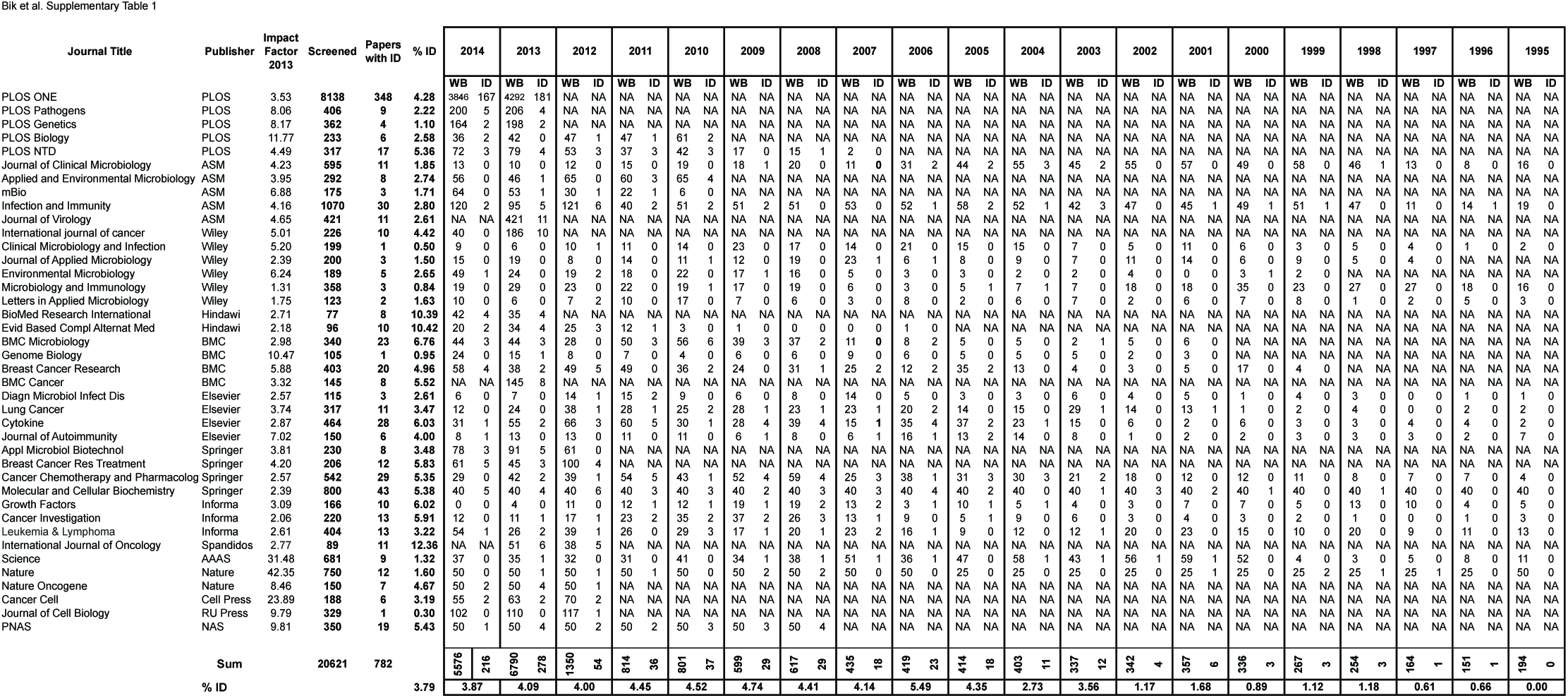
The 40 journals screened in this study. Table includes publisher, impact factor (Thomson Reuters 2013), number of papers containing the term “western blot” (WB) screened per year, and number of papers with inappropriate image duplication (IDs) found in that year.

